# The Developmental Trajectory of the Social Brain: A Movie-Based Exploration from childhood to adolescence

**DOI:** 10.1101/2025.07.24.666623

**Authors:** Sara Saljoughi, Taylor Heffer, Mehran Ebrahimi, Kathleen Lyons, Bobby Stojanoski

**Affiliations:** Faculty of Science, Ontario Tech University, Oshawa, ON; Faculty of Social Science and Humanities, Ontario Tech University, Oshawa, ON; Department of Psychology, King’s University College, London, ON; Department of Psychology, Western University, ON, Canada; Western Institute for Neuroscience, Western University, ON, Canada

**Keywords:** Social cognition, Affective and cognitive Theory of Mind, Empathy, Development, Childhood, Adolescence

## Abstract

In childhood and adolescence, functional brain networks go through different stages of development, and the levels of connectivity within and between these networks change with age. The developmental trajectory of these large-scale networks of the brain have been extensively investigated; however, many aspects of our social brain and its developmental patterns remain unclear. This study employed a cross-sectional design to investigate the brains of 753 children and adolescents (ages 5-15) while they watched a movie. This research investigates the functional distinctness and developmental synchronicity of brain areas implicated in social cognition, such as empathy and affective and cognitive theory of mind, across childhood and adolescence using generalized additive models and inter-region group analysis. Our findings suggest that social cognition components networks such as cognitive and affective theory of mind and empathy exhibit distinct developmental trajectories throughout childhood and adolescence. The findings support the theory that social cognitive networks are developmentally distinct from each other, even in the absence of task-specific paradigms.

## Introduction

As children mature into adolescents, they experience profound changes to all aspects of their social lives. During this time, children’s social networks begin to grow, and their interactions with others become more structured and complex (Adolphs, 2009; Casey et al., 2008; Crone & Dahl, 2012; Dunbar, 1998). Expanding social lives during this period is underscored by considerable change to various aspects of social cognition, such as, theory of mind (ToM), which is the ability to infer others’ beliefs, intentions, and desires(Frith & Frith, 2007), and empathy, which is the capacity to grasp and share others’ affective state(Schurz et al., 2021a). These social cognitive abilities reflect an individual’s ability to make sense of social environments, form the basis of children’s social lives, and are central to the way children think, interact, and live in social contexts (Blakemore, 2008; Blakemore & Mills, 2014).

Not coincidentally, this developmental epoch coincides with significant maturation of underlying neural circuitry associated with various aspects of social cognition. Specifically, the “social brain”(Adolphs, 2009; Blakemore, 2008; Casey et al., 2008; Crone & Dahl, 2012; Dunbar, 1998; Frith & Frith, 2007), is comprised of a distributed network of brain regions including the medial prefrontal cortex (mPFC), temporoparietal junction (TPJ), posterior superior temporal sulcus (pSTS), amygdala, and precuneus(Adolphs, 2009). This network supports the diverse set of processes associated with theory of mind and empathy, such as interpreting social signals generated by others (Frith & Frith, 2021; Lieberman, 2007; Magno et al., 2022; Saxe & Kanwisher, 2003), understanding the mental state of others, and mirroring others emotions(Adolphs, 2009). Although the social brain is often represented as a single functional network, emerging early in life and remaining relatively unchanged after late childhood, there is evidence to suggest the social brain contains distinct subnetworks that are associated with different aspects of social cognition (McCormick et al., 2018). For instance, the social brain can be broken down into specific networks for theory of mind including the medial prefrontal cortex (MFPC), precuneus (PC), and bilateral temporoparietal junction (RTPJ/LTPJ) and for empathy-related processes like pain perception, which involve regions such as right and left insula, right and left medial frontal gyrus (MFG) and the dorsal anterior middle cingulate cortex(Richardson et al., 2018). These networks are functionally distinct as early as three years of age and continue to differentiate throughout development into adulthood(Richardson et al., 2018). Indeed, theory of mind and empathy are associated with developmentally distinct functional neural profiles as well (Shamay-Tsoory et al., 2009; Shamay-Tsoory & Aharon-Peretz, 2007). In fact, even the theory of mind network can be decomposed into subnetworks reflecting distinct affective and cognitive components: cognitive ToM refers to the ability to infer others’ beliefs, intentions, and knowledge, and affective ToM involves making inferences about others’ emotional states (Arioli et al., 2021a; Dunbar, 1998; Richardson et al., 2018; Schurz et al., 2021b). A similar division may also be true for empathy (Molenberghs et al., 2016), whereby affective empathy refers to the emotional sharing or resonance with others’ feelings, and cognitive empathy involves understanding others’ emotions through reasoning (Molenberghs et al., 2016). Notably, affective empathy is commonly referred to simply as “empathy”, and cognitive empathy closely aligns with affective ToM, as both involve reasoning about others’ emotions(Shamay-Tsoory et al., 2010). This conceptualization aligns with a recent proposed hierarchical model in which social cognitive processes are organized into three clusters: a cognitive cluster (reflecting cognitive ToM), an affective cluster (representing affective empathy), and an intermediate cluster (capturing the affective ToM) (Arioli et al., 2021a; Schurz et al., 2021b). Taken together, these three components (cognitive ToM, affective ToM, and empathy) can be considered as the core building blocks of our social brain (Arioli et al., 2021b; Schurz et al., 2021a; Shamay-Tsoory et al., 2010). These set of findings suggest that the social brain contains functionally and developmentally distinct subnetworks associated with different aspects of social cognition (Schurz et al., 2021b).

Traditionally, investigating the development of different socio-cognitive systems supported by regions of the social brain have relied on task-based fMRI, using highly controlled stimuli to probe specific functions (e.g., viewing faces, theory-of-mind vignettes)(Fu et al., 2023). While invaluable, these approaches have limitations: task-based paradigms often employ simple stimuli that may not capture the complexity of assessing the neural mechanisms of aspects of social cognition (Fu et al., 2023; Quesque & Rossetti, 2020). As a result, many of the tasks and paradigms used to assess the neural mechanisms of social cognitive abilities, like theory of mind and empathy, yield conflicting results(Quesque & Rossetti, 2020). More recently, task-free paradigms, such as resting-state fMRI, have revealed fundamental aspects of intrinsic network organizations, and the changing developmental trajectories of the functional connectivity architecture of these networks(Sanders et al., 2023; Somerville et al., 2018). However, much like task-based approaches, resting-state fMRI fail to capture the richness and dynamism of processing real-world social situations(Sonkusare et al., 2019). Bridging the gap between controlled experimental settings and the complexity of real-world social cognition requires paradigms that offer more ecological validity (Sonkusare et al., 2019).

Movie watching provides a means of bridging this gap and is ideal for examining the development of the social brain(Vanderwal et al., 2019). Movies provide complex, dynamic, and continuous streams of information that closely mimic the perceptual and cognitive demands of real-world social observation(Lyons et al., 2020a; Sonkusare et al., 2019). They contain rich social narratives, character interactions, emotional expressions, and implicit mental states that naturally engage the social brain network(Lyons et al., 2020a; Saarimäki, 2021) and induce a more uniform social cognitive state across participants compared to other task-free states such as resting-state and non-plot based movies (Hasson et al., 2004; Vanderwal et al., 2015). By examining brain activity as participants process the social narrative, movie-watching fMRI allows us to investigate how neural systems respond to and interpret complex, life-like social information. Although movie-watching has emerged as a powerful tool for examining changes to large scale network architecture across development(Finn et al., 2015; Hasson et al., 2004; Lyons et al., 2020a; Sonkusare et al., 2019; Vanderwal et al., 2019; Yu, 2024), distinct and overlapping developmental trends of the social brain in response to social aspects of movies has not yet been examined.

In the current study, we combine fMRI movie-watching and Generalized Additive Models (GAMs) to identify whether subnetworks of the social brain are best captured by linear or non-linear developmental trajectories (Wood, 2017). We focused on three components (cognitive and affective theory of mind and empathy) as the core building blocks of the social brain. GAMs provide a powerful and flexible statistical framework for characterizing age-related changes in functional connectivity profiles within and between subnetworks of the social brain, potentially revealing accelerating, decelerating, or plateauing developmental windows (Choi et al., 2023; Sanders et al., 2022; Tooley et al., 2022) in response to socially rich content. This is particularly important during the dynamic periods of brain development in childhood and adolescence which do not always follow linear trends (Gogtay et al., 2004; McCormick et al., 2018; Sanders et al., 2022; Shaw et al., 2008), and assuming linearity risks obscuring important developmental periods of change (Crone & Dahl, 2012; Fjell et al., 2012; Luna et al., 2015). Based on previous work, we hypothesized that functional connectivity within the social brain network during movie watching will show significant age-related changes, reflecting increased specialization supporting increasingly sophisticated social processing abilities (Crone & Dahl, 2012; Saxe & Kanwisher, 2003). We also predicted that these developmental trajectories for the different subnetworks comprising the social brain will follow distinct developmental trajectories following both linear and non-linear trends. Identifying distinct developmental trends within each sub-network present the possibility that these regions may be functionally distinct and contribute to different aspects of social cognition.

## Materials and methods

### Participants

A total of 780 children and adolescent data sets from the Healthy Brain Network (HBN) biobank (releases 1 to 8) (Alexander et al., 2017), an initiative of the Child Mind Institute were included in the study. All data were previously preprocessed (Lyons et al., 2020b, 2024; Pho et al., 2024), and passed quality assurance based on visual inspection of T1-weighted images and evaluation of functional connectivity matrices to ensure data integrity. Of the 780, 27 participants were excluded for excessive motion (greater than 1.5 mm). The final data set included 753 children and adolescents aged 5-15 years (mean=10.19, std=2.57; 249 female). Phenotypic data included age and sex assigned at birth (Table 1 presents a summary of participant demographics). Participants were not excluded based on clinical diagnoses to ensure the sample reflect a representative sample of the population. The Chesapeake Institutional Review Board approved the study, and details on the HBN biobank can be found here: http://fcon_1000.projects.nitrc.org/indi/cmi_healthy_brain_network/. The institutional review board at Ontario Tech University approved secondary analysis of the HBN data.

**Table 1.**
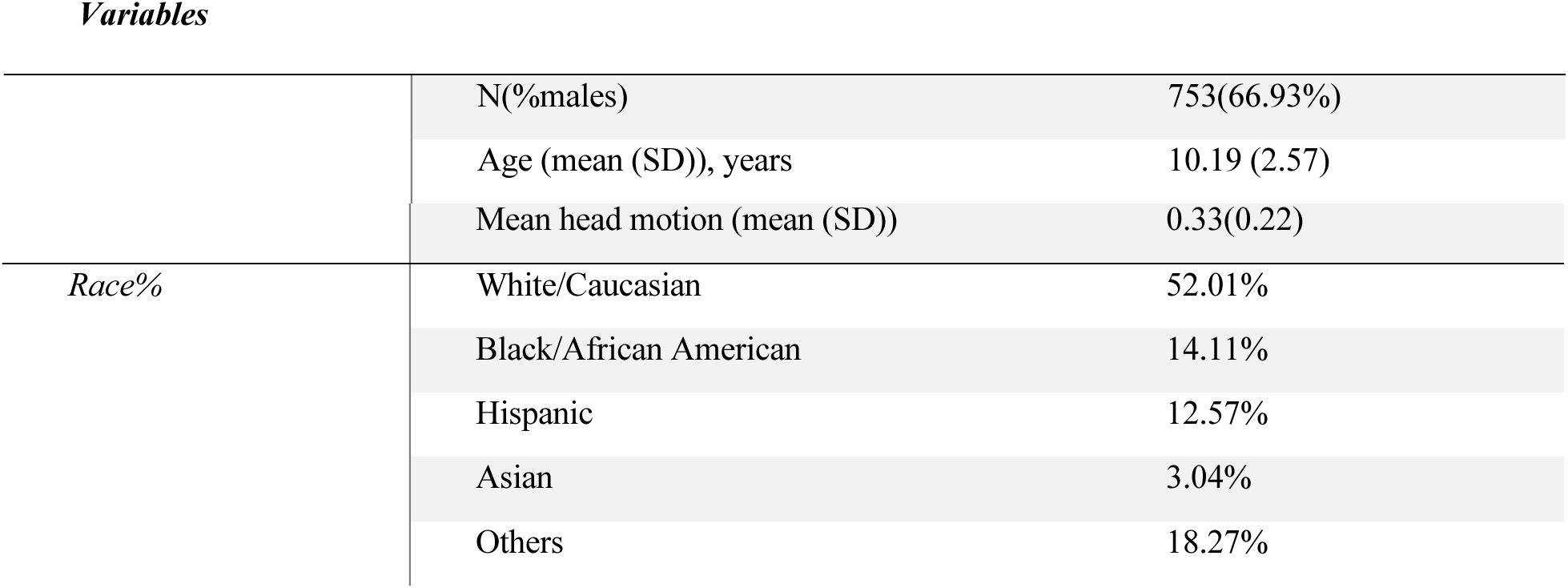
Summary of participant demographics.

*Variables*
|  |  |
| --- | --- |
| N(%males) | 753(66.93%) |
| Age (mean (SD)), years | 10.19 (2.57) |

Table1- Summary of participant demographics.
|  |  |  |
| --- | --- | --- |
|  | Mean head motion (mean (SD)) | 0.33(0.22) |
| *Race%* | White/Caucasian | 52.01% |
|  | Black/African American | 14.11% |
|  | Hispanic | 12.57% |
|  | Asian | 3.04% |
|  | Others | 18.27% |

### fMRI stimuli

Participants underwent T1-weighted anatomical and functional magnetic resonance imaging (MRI) during a 10-minute clip from the animated film ’Despicable Me’. We chose to use data from this scan because movies can be powerful tools for studying social cognition in the brain. In particular, ’Despicable Me’ requires careful processing of the plot, character development and contains rich social interactions which mimic real-world experiences making it ideal for investigating the underlying brain mechanisms associated with social cognitive processes (Vanderwal et al., 2019).

### fMRI acquisition and processing

MRI data was acquired using a 3 Tesla Siemens scanner with a 32-channel head coil. Structural scans were obtained using a sagittal sequence (TR=2500 ms, TE=3.15 ms, resolution=0.8 mm isotropic) and functional scans used a gradient-echo planar imaging sequence (TR=800 ms, TE=30 ms, Flip Angle=31 degrees, whole brain coverage with 60 slices, resolution=2.4 mm isotropic).

The resulting neuroimaging data were preprocessed and analyzed with the Automatic Analysis (AA) toolbox, SPM 8, and in-house MATLAB scripts. Functional data preprocessing involved motion correction utilizing six motion parameters (translation: left/right, anterior/posterior, superior/inferior; rotation: pitch, roll, yaw). Finally, both functional and structural images were normalized to a standard brain template (Montreal Neurological Institute, MNI) to enable consistent analysis across participants. Following a spatial smoothing of the functional data with a Gaussian filter (8 mm kernel), drift and other low-frequency noise were eliminated by high-pass filtering with a threshold of 1/128 Hz. To remove unwanted noise from the data, we applied bandpass filtering. This technique identifies and removes signals from the cerebrospinal fluid, white matter, head movements, and sudden spikes in the signal as noise.

Functional connectivity matrices were computed for every participant, based on 49 brain seed regions which together encompass the social brain, including cognitive ToM, affective ToM, and empathy (see Figure 1). These regions were defined based on a large-scale meta-analysis that identified consistent activation patterns within the social brain, independent of input sensory modality, stimulus type, or cognitive task (Arioli et al., 2021a). Regions of interest (ROIs) were defined using MRIcron software based on the spatial coordinates outlined. Functional connectivity within these regions was then calculated with Pearson correlations between the activity time series of each ROI pair. The resulting correlation coefficients were standardized using z-scores and compiled into a 49 x 49 functional connectivity matrix for each participant.

**Figure 1.**
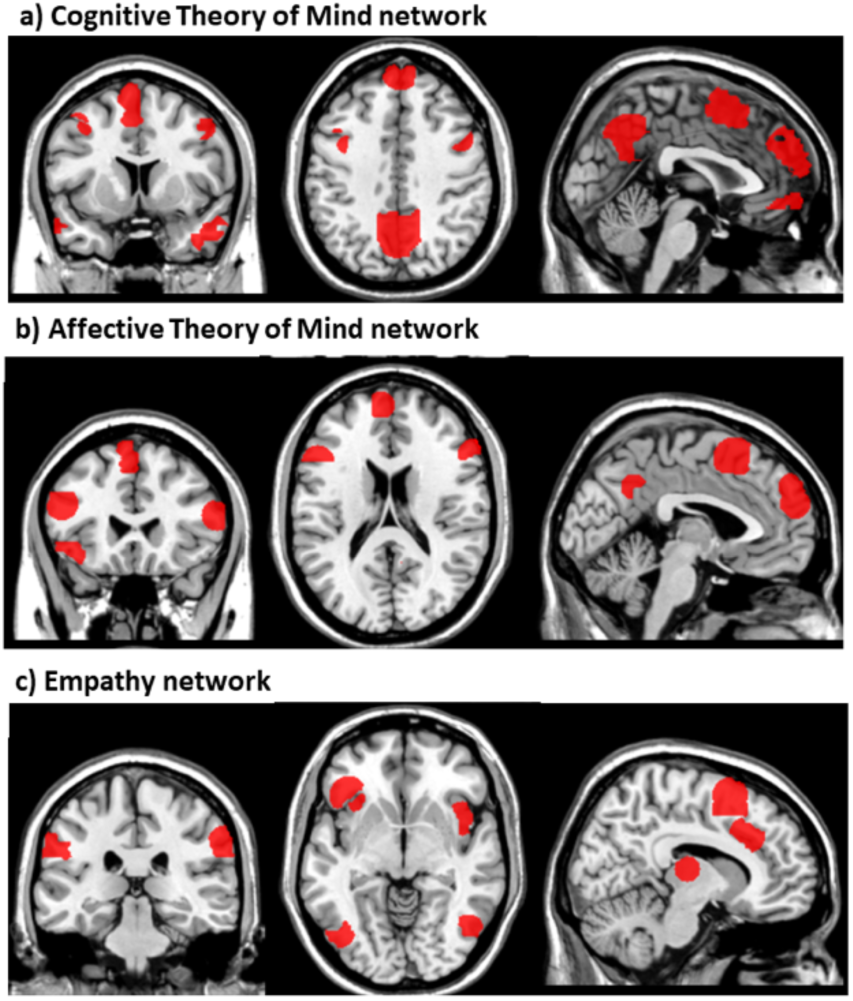
Regions of Interest (ROIs) for Social Brain networks: (a) Cognitive ToM, (b) Affective ToM, and (c) Empathy.

### Analytic approaches

We employed two complementary approaches to investigate potential changes in functional connectivity within and between social brain networks across development. First, generalized additive models (GAMs) were used to explore linear or non-linear developmental patterns across the different social brain networks. Second, group contrasts were employed to identify connectivity profiles across brain regions (nodes within and between each network) that underwent developmentally meaningful changes.

### GAMs

We used generalized additive models (GAMs) to analyze age-related differences in both within- and between-network connectivity across three social brain networks: affective ToM, cognitive ToM, and empathy. GAMs utilize cross-validation to prevent overfitting and can capture complex, non-linear relationships between age and functional connectivity. In contrast with traditional models that are limited to linear or polynomial functions, GAMs produce data-driven, smooth curves that reflect the underlying patterns (Wood, 2017). In GAMs, ’effective degrees of freedom’ (EDF) reflects how complex a relationship is between variables. An EDF of 1 indicates a linear relationship, whereas larger EDF values represent more complex non-linear relationships (Wood, 2017). GAM models were fit with Restricted maximum likelihood (REML) to mitigate over-fitting (Wood, 2001, 2017).

We employed generalized additive models (GAMs) implemented in the R package ’mgcv’ (version 1.8.42) (Wood, 2017) within R software (version 4.3.1; R Core Team, 2019) and scatterplots were created using the ggplot2 package (Wickham, 2016) in R.

We investigated the relationship between age and functional connectivity within and between the three social brain networks, functional connectivity was computed by averaging Fisher Z-transformed pairwise correlations within and between nodes in each of the social brain networks. As additional factors to account for their possible impact on the results, sex and mean framewise displacement (FD (Power et al., 2012): age-related differences in in-scanner motion are well established; children have a greater tendency to move their heads during scan sessions (Satterthwaite et al., 2019), which can impact measures of functional connectivity (Power et al., 2013; Van Dijk et al., 2012)), were included as covariates. To visualize and interpret how age interacts with the observed trend, shaded regions corresponding to standard error of the mean were plotted around the curve.

### Inter-region group analyses

Motivated by the observed heterogeneity in the developmental trajectories of the social brain networks, we sought to identify the precise brain areas (nodes) which may be most strongly linked to the developmental change in functional connectivity profiles. To investigate developmental trends, we divided participants into four non-overlapping age groups: early childhood (5–7.5 years, *n* = 122), middle/late childhood (7.51–10 years, *n* = 267), early adolescence (10.01–12.5 years, *n* =197), and middle/late adolescence (12.51–15.5 years, *n* = 168). Group comparisons were conducted using ANCOVA on the functional connectivity values for each edge, controlling for head motion (FD) as a covariate. We used a modified F-statistic that incorporates the sign of the mean difference between groups (for example, the direction for late childhood > early childhood) enabling interpretation of both the effect size and its directionality. The significance threshold was set at α = 0.05, and multiple comparisons were corrected using the False Discovery Rate (FDR) method.

## Results

### Age-related differences in sub-social networks

Age-related changes in within- and between-network connections were assessed for the three social brain networks using GAMs (Fig 2). We found six distinct age-related patterns, all statistically significant. That is, the cognitive ToM network showed a non-linear increase across age best represented with a quadratic-like pattern (edf = 2.416, ref.df = 3.028, F = 3.630, p = 0.0124; see Fig. 2). In contrast, the results of the GAM for the affective ToM network revealed a linear relationship (edf = 1.001, ref.df = 1.002, F = 6.356, p = 0.01), and the GAM results for empathy networks suggest a more complex, cubic-like pattern (edf = 2.971, ref.df = 3.714, F = 2.503, p = 0.0406). We also examined the between network connections. We found that functional connectivity between affective and cognitive ToM (edf = 1.758, ref.df = 2.202, F = 3.989, p = 0.0167), affective ToM and empathy (edf = 4.402, ref.df = 5.428, F = 3.133, p = 0.0068) and cognitive ToM and empathy (edf = 3.174, ref.df = 3.962, F = 3.159, p = 0.0143) all showed a non-linear increase across age, which may reflect a potential sensitive period of the social brain during development.

**Figure 2.**
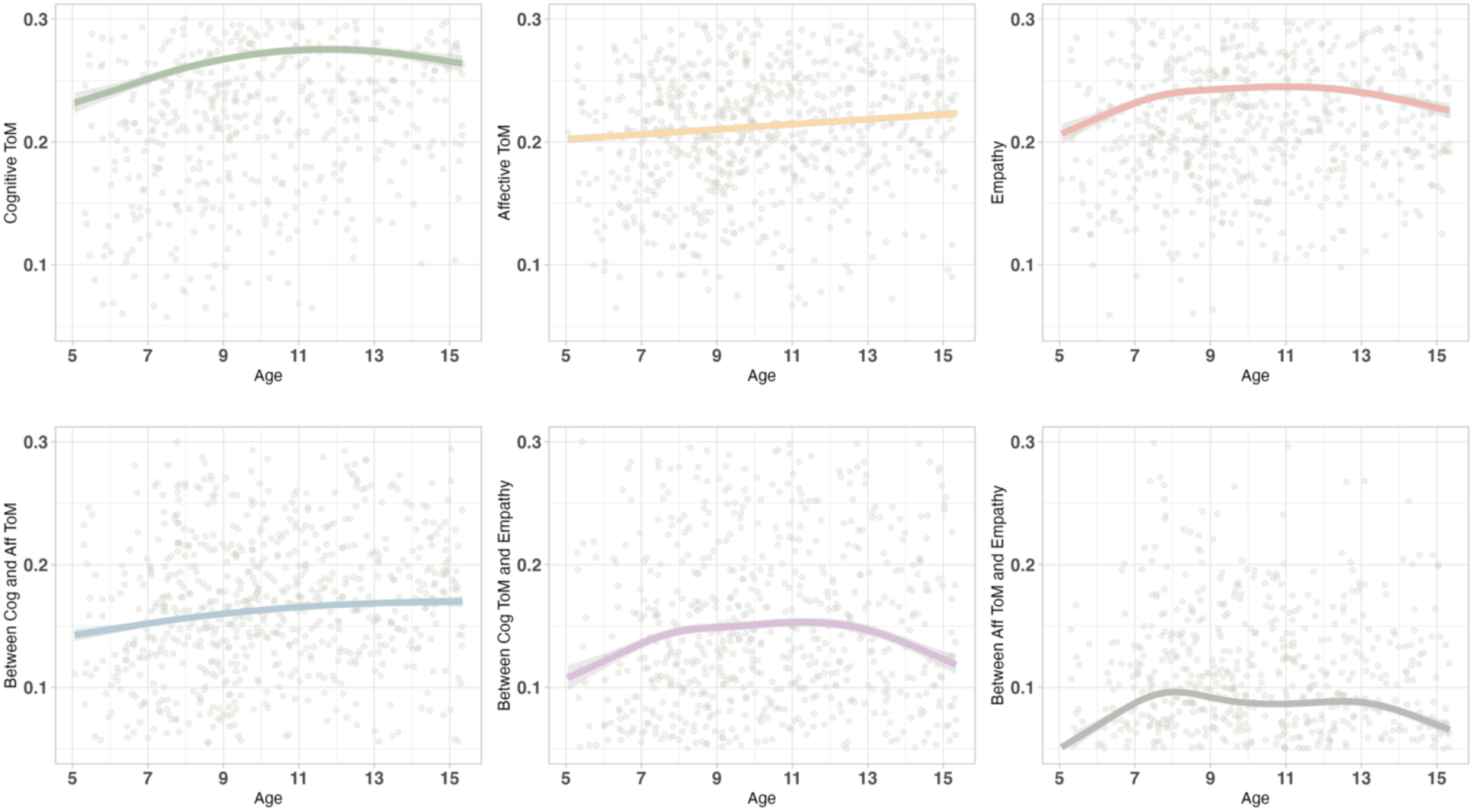
Developmental trajectories of functional connectivity within and between social brain networks-Generalized Additive Models (GAMs) were used to model age-related changes in functional connectivity across the three social brain networks. Top row: Within-network connectivity for (left) cognitive theory of mind, (middle) affective theory of mind, and (right) empathy. Bottom row: Between-network connectivity for (left to right) cognitive– affective ToM, cognitive ToM–empathy, and affective ToM–empathy. Shaded areas represent standard error of the mean.

### Inter-region correlation analysis

The inter-region correlation analysis was employed on four groups: early childhood, middle/late childhood, early adolescence, and middle/late adolescence. Figure 3 illustrates the significant connections exhibiting changes during development. We compared the earliest group with the three subsequent age groups. Our findings revealed an overall trend of strengthening connections between brain components as participants aged. Notably, panel (a) of Figure 3 reveals a general trend of strengthening within-network connectivity across the three social brain networks (cognitive and affective ToM, and empathy) over development. Specifically, we found an increase in the number of significantly connected nodes within social brain networks. When comparing the earliest age group (early childhood group) to the middle/late childhood group, there was a marked increase of 187 significant connections. Further analysis revealed 196 significant connections between the first and third age groups, and a total of 363 more significant connections in the oldest group compared to the youngest. We also examined which regions showed the strongest developmental changes in connectivity both within and between subnetworks of the social brain (represented by the largest absolute F-statistic with values exceeding 10). Interestingly, our results suggest segregation of the cognitive ToM network, with a distinct cluster forming between temporal regions that is present in the adolescent group but not present at early and middle/late childhood (as shown in Figure 3a). Also, in the comparison between early childhood and middle/late adolescence, a set of negative connections within the empathy network, primarily involving the left inferior occipital gyrus and the right fusiform gyrus (Figure 3b) emerged, which were not present in the younger groups (Figure 3a, b). In contrast, the analysis of between-network connectivity suggests a trend towards increased negative connections between the social brain networks, with the strongest effects showing modified F-statistics lower than -10, as illustrated in Figure 3 c. That is, we found more negative connections (correlations) between the brain networks, particularly between the empathy network and both the cognitive and the affective ToM networks, across development, with the most negative connections in the adolescent group. Notably, the comparison between the oldest group compared to the youngest revealed several prominent positive connections, including: connectivity between 1) Right superior temporal gyrus and right middle temporal gyrus (F = 53.23, p<0.001), 2) Left middle temporal gyrus and left superior temporal gyrus (F = 52.10,p<0.001), 3) Right inferior temporal gyrus and right middle temporal gyrus (F = 50.42, p<0.001), 4) Left superior temporal gyrus and right middle temporal gyrus (F = 47.32, p<0.001), and 5) Left superior temporal gyrus and right middle temporal gyrus (F = 45.89, p<0.001). Additionally, several strong negative connections were observed including 1) Left insula and right middle temporal gyrus (F = 34.74, p<0.001), 2) Left inferior occipital gyrus and middle cingulate gyrus (F = 33.82, p <0.001), and 3) Left inferior occipital gyrus and right supplementary motor area (F = 26.54, p <0.001).

**Figure 3.**
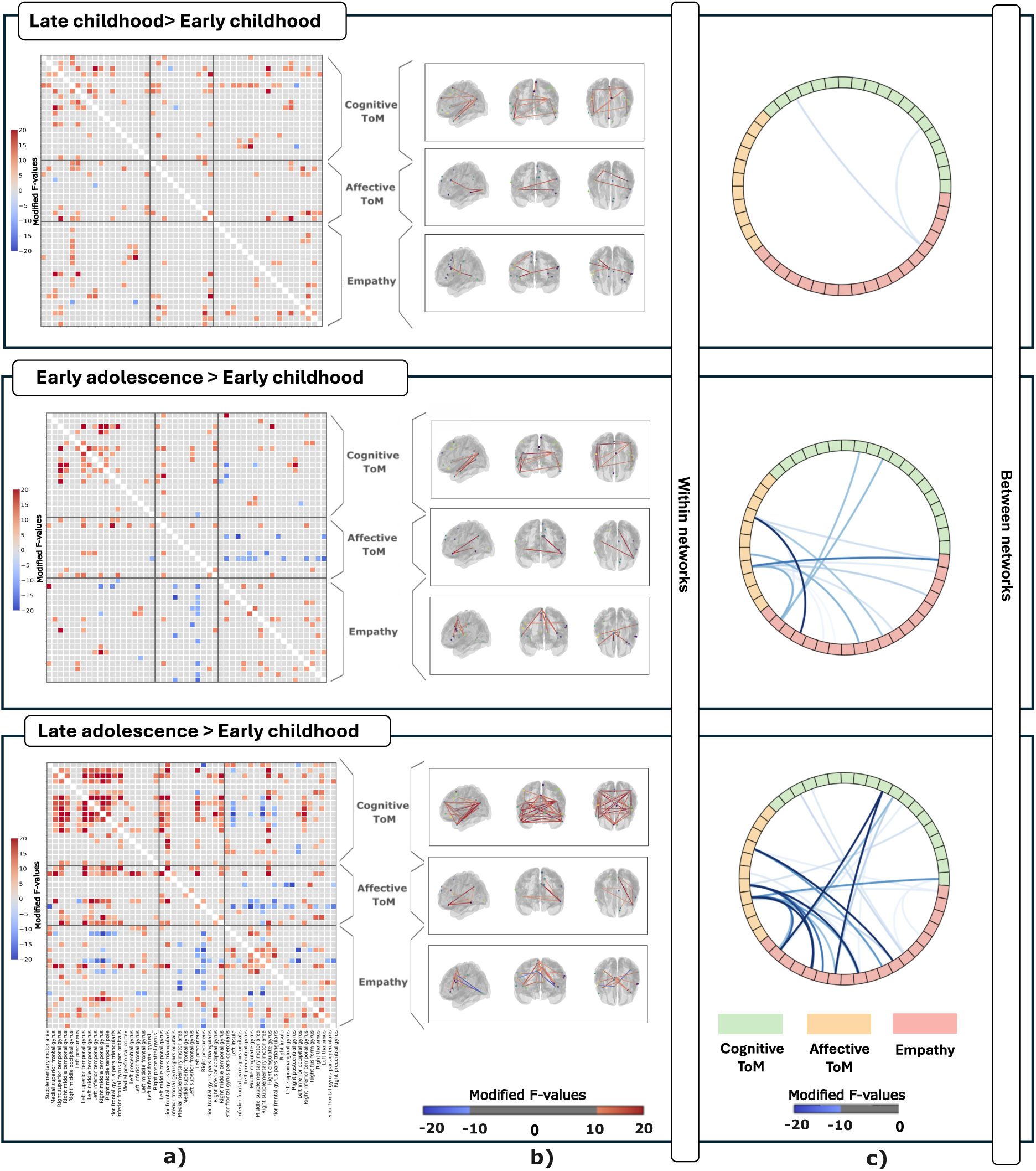
Inter-region correlation analysis across age bins reveals developmental changes in connectivity within and between social brain networks. a) FDR-corrected ANCOVA comparisons of inter-region connectivity between early childhood and each subsequent group, controlling for head motion (FD): middle/late childhood, early adolescence, and middle/late adolescence, Red and blue squares denote increases and decreases in connectivity, respectively, based on a modified F-statistic, calculated by multiplying the F-value by the difference in means between the two groups, b), Strongest developmental effects in within-network connectivity (|modified F-values | < 10), with rows showing: (1) cognitive ToM, (2) affective ToM, and (3) empathy networks, corresponding to the same pairwise comparisons. c) Significant negative connectivity between subnetworks of the social brain networks across development, based on the same pairwise comparisons.

## Discussion

The current study used a cross-sectional movie-watching fMRI paradigm to examine age-related developmental changes in functional connectivity both within and between subnetworks of the social brain from childhood to adolescence. We found distinct linear and non-linear age-related developmental trajectories for each of the three subnetworks comprising the social brain. Both linear and non-linear age-related changes were driven by a combination of increasing positive within network connections and negative between network connections. Although we found increases in positive correlations across development in all three subnetworks, these changes were strongest during middle/late adolescence and primarily within the cognitive theory of mind network. Similarly, the most and strongest increase in negative correlations were found during middle/late adolescence and these changes were widespread between all three subnetworks.

We found distinct developmental trajectories in within-network connectivity across the social brain subnetworks. The cognitive theory of mind network demonstrated a rapid increase during early development, followed by a plateau and consistently, reached the highest overall connectivity values across all six comparisons. The affective ToM network showed a linear increase across age. In contrast, the empathy network exhibited an inverted U-shaped trajectory, with connectivity increasing during mid-childhood and decreasing in adolescence. Each of these sub-networks are associated with distinct yet interrelated cognitive functions. Cognitive ToM involves reasoning about others’ thoughts, motivations, and beliefs (Arioli et al., 2021b; Schurz et al., 2014) and draws heavily on executive functions (Schurz et al., 2014). Affective ToM, on the other hand, supports inferences about what others are feeling (Shamay-Tsoory et al., 2010). The empathy network also supports the capacity to share and understand others’ emotional and sensory experiences (Mercadillo et al., 2011; Xiong et al., 2019). These results suggest that despite forming a singular “social brain”, these subnetworks are developmentally distinct, likely reflecting the associations with the maturing of specific social cognitive abilities. Indeed, our findings align with previous work distinguishing between cognitive and affective components of ToM and empathy (Shamay-Tsoory et al., 2010).

Additional evidence for distinct developmental trajectories of the subnetworks forming the social brain emerges from our results examining between subnetwork connectivity. We found that all between-network connections, that is, between cognitive-affective, cognitive-empathy, and affective-empathy networks, also showed significant nonlinear patterns across age, suggesting more complex, stage-specific developmental shifts. These age-related changes in connectivity likely reflect meaningful developmental differences in the integration and coordination of cognitive and affective processes shaped by social experiences from childhood through adolescence. Notably, the developmental pattern of connectivity between cognitive and affective theory of mind may signal continued integration and refinement of the networks most directly involved in higher-order social cognitive functions, such as understanding others’ beliefs, intentions, and emotions—abilities known to undergo prolonged development into late adolescence (Blakemore, 2012). These findings suggest that subregion-specific processing of affective and cognitive information may each contribute in a graded manner to a broader, unified ToM function (Schurz et al., 2014). In contrast, connections between the empathy network and both ToM networks show a developmental trend toward increased segregation. The overlapping yet developmentally distinct neural recruitment for cognitive and affective theory of mind and empathy suggests their co-development and increasing interaction across childhood and adolescence (Richardson et al., 2018; Sebastian et al., 2012). Age-related changes in connectivity between subnetworks likely reflect developmentally meaningful differences in the efficacy with which specific information is processed and are shaped by experiences from childhood to adolescence.

Consistent across all analyses involving the empathy network (both within-network connectivity and its connections with cognitive and affective theory of mind networks) was a pattern of increasing connectivity in early childhood, followed by a plateau in early adolescence and a decline in middle to late adolescence. This trajectory may reflect the early maturation of foundational affective sharing mechanisms. Early in life, widespread empathic connectivity reflects the brain’s immature and expansive architecture designed to capture affective experiences (Decety, 2010). During early adolescence, this connectivity plateaus, marking a transitional phase where empathy becomes finely tuned, and aligns with recent evidence that empathy matures considerably during early adolescence, reflecting ongoing refinement and restructuring of the brain’s social systems (Gaspar & Esteves, 2022). Finally, the decline seen in middle and late adolescence aligns with synaptic pruning and neuroplasticity, as the brain streamlines redundant connections and enhances processing efficiency, yielding more specialized empathy circuits as adolescents prepare for adult social functioning (Konrad et al., 2013).

Across all three networks, we observed a developmental increase in the strength and number of nodal connections within sub-networks across development, that were strongest in middle and late adolescence, which suggests that these areas become more functionally cohesive, facilitating improved processing of social information. This maturation is thought to underpin the enhanced ability of adolescents to navigate complex social environments, interpret nuanced social cues, and engage in sophisticated perspective-taking.

We found significant increases in the amount and strength of connections within the temporal compared to the frontal nodes of the cognitive ToM network. This is important because the hierarchy of social inferences is based on the continuous information provided by lower-level brain areas (such as the amygdala) to higher order regions, such as TPJ/pSTS to guide trait interpretations of the agent (Frith & Frith, 2021). Although prior research has emphasized the importance of frontal regions in cognitive functions like theory of mind (Amodio & Frith, 2006; Frith & Frith, 2021), recent studies highlight the crucial role of temporal regions in similar processes. For instance, key temporal nodes, including the temporoparietal junction (TPJ), posterior superior temporal sulcus (pSTS), and middle temporal gyrus (MTG), are crucial for processing complex dynamic social cues such as gaze, motion, goal-directed behavior, and even mind-reading during social interactions (Braunsdorf et al., 2021; Mars et al., 2012). These mid-level brain areas act as intermediaries in a social inference hierarchy, integrating affective input from lower-level regions (e.g., the amygdala) and providing interpreted social signals to higher-order regions such as the mPFC, which supports trait-based inferences and mental state reasoning (Frith & Frith, 2021). These findings suggest that temporal regions might be a key factor in our unique social abilities, differentiating us from other species in the context of theory of mind (Frith & Frith, 2021; Zaki & Ochsner, 2012). This increased connectivity allows for more integrated information processing within the temporal cortex itself, and between this region and other brain areas involved in higher-order functions. This enhanced communication within the network likely underlies the complex behaviors that distinguish humans from other species (Braunsdorf et al., 2021). On the other hand, the stronger temporal connectivity we observed may also indicate a developmental shift toward greater efficiency and automation in social cognitive processing. As theory of mind abilities becomes more practiced and embedded in everyday functioning, the brain may increasingly rely on lower-energy, faster temporal circuits for routine social understanding, reducing dependence on resource-intensive frontal regions (Dumontheil, 2016). Altogether, these findings suggest the temporal cortex’s central role in supporting both foundational and increasingly efficient cognitive ToM abilities during development.

Our results also revealed a developmental increase in negative correlations between these social brain networks, particularly between the empathy network and both cognitive and affective ToM networks. This increase in negative correlations, specifically between connections between the empathy network and both ToM networks, provide support for a trend toward segregation. In parallel, the emergence of negative connections within the empathy network itself during middle/late adolescence may reflect similar underlying mechanisms, such as synaptic pruning or increasing intra-network segregation, potentially signaling continued developmental reorganization. This pattern suggests the emergence of distinct subcomponents within the social brain, potentially reflecting functional specialization of these networks in supporting different aspects of social cognition. Such segregation may be a marker of increasing neural efficiency, where functional systems reduce redundant interactions and optimize information processing for distinct cognitive tasks (Achard & Bullmore, 2007; Cherniak, 1994). As social cognitive abilities such as cognitive ToM, affective ToM, and empathy become more differentiated, their underlying neural mechanisms may become increasingly distinct, supporting more specialized forms of mental state reasoning and emotional responsiveness. Taken together, these developmental changes suggest a reorganization of the social brain in development, with refined boundaries between subsystems that allow for greater flexibility and context sensitivity in social cognition.

## Limitations

Our results are based on children and adolescents 5 years of age and older, however, early childhood is a critical period for brain development. Humans are born with a proclivity for social connection and rely on their group for survival driving the development and learning of social cognitive skills that act as "wiring instructions" that shape the developing social brain (Atzil et al., 2018). For a more comprehensive understanding of the development of the social brain, it would be important for future research include younger participants. In addition, this work is cross-sectional. Although our results have shed light on the differential developmental trajectories of the different subnetworks of the social brain, future work using longitudinal paradigms to examine variability from a within participant perspective is important.

## Conclusion

Adolescence and childhood are critical stages characterized by significant cognitive, social, and behavioral development. Given the continuous nature of brain development, identifying developmental milestones is challenging. To capture this gradual change, we employed two complementary approaches. First, we utilized GAMs to analyze the overall trends in functional connectivity across the age range of interest. Second, we employed an analysis based on age bin groups to pinpoint specific brain regions that exhibit the most pronounced changes in connectivity. Examining the specific the developmental changes to social brain is not only important for understanding how children respond to a changing social landscape, but may have important implications for both neurotypical and neurodivergent development, as social challenges are commonly implicated in a range of neurodevelopmental conditions and may reflect underlying differences in the development of the social brain (Blakemore, 2008; Pelphrey et al., 2011).

## Supporting information

The labels and brain regions related to cognitive ToM, affective ToM, and empathy

## Acknowledgements

We would like to thank the Child and Mind Institute for designing and collecting data for the Healthy Brain Network Biobank. We would also like to thank the children, adolescents, and their families for taking the time to participate in studies conducted by the Child and Mind Institute. This work was funded by a Natural Sciences and Engineering Research Council of Canada Discovery grant (RGPIN-2020-05042 to BS)

