## Supplementary material for "The Developmental Trajectory of the Social Brain: A Movie-Based Exploration from childhood to adolescence": The labels and brain regions related to cognitive ToM, affective ToM, and empathy

**Supplementary document:**

**The labels and brain regions related to cognitive ToM, affective ToM, and empathy used in this paper(Arioli et al., 2021b)**

|  | **brain regions** |
| --- | --- |
| **Cognitive ToM** | **'Supplementary motor area'** |
|  | **'Medial superior frontal gyrus'** |
|  | **'Right superior temporal gyrus'** |
|  | **'Right middle temporal gyrus'** |
|  | **'Right middle occipital gyrus'** |
|  | **'Left precuneus'** |
|  | **'Left superior temporal gyrus'** |
|  | **'Left middle temporal gyrus'** |
|  | **'Left inferior temporal gyrus'** |
|  | **'Right middle temporal gyrus'** |
|  | **'Right middle temporal pole'** |
|  | **'Right inferior frontal gyrus pars triangularis'** |
|  | **'Right inferior frontal gyrus pars orbitalis'** |
|  | **'Medial prefrontal cortex'** |
|  | **'Left precentral gyrus'** |
|  | **'Left inferior frontal gyrus'** |
|  | **'Left middle frontal gyrus'** |
|  | **'Left inferior frontal gyrus1_'** |
|  | **'Right precentral gyrus_'** |
| **Affective ToM** | **Left middle temporal gyrus'** |
|  | **'Left inferior frontal gyrus pars triangularis'** |
|  | **'Left inferior frontal gyrus pars orbitalis'** |
|  | **'Medial supplementary motor area'** |
|  | **'Medial superior frontal gyrus'** |
|  | **'Left superior frontal gyrus'** |
|  | **'Left precuneus'** |
|  | **'Right precuneus'** |
|  | **'Right inferior frontal gyrus pars triangularis'** |
|  | **'Right inferior occipital gyrus'** |
|  | **'Right middle temporal gyrus** |
| **Empathy** | **Left inferior frontal gyrus pars opercularis'** |
|  | **'Left insula'** |
|  | **'Left inferior frontal gyrus pars orbitalis'** |
|  | **'Left precentral gyrus'** |
|  | **'Middle cingulate gyrus'** |
|  | **'Middle supplementary motor area'** |
|  | **'Right supplementary motor area'** |
|  | **'Right cingulate gyrus'** |
|  | **'Right inferior frontal gyrus pars triangularis'** |
|  | **'Right insula'** |
|  | **'Left supramarginal gyrus'** |
|  | **'Right postcentral gyrus'** |
|  | **'Left inferior occipital gyrus'** |
|  | **'Right inferior temporal gyrus'** |
|  | **'Right fusiform gyrus'** |
|  | **'Right thalamus'** |
|  | **'Left thalamus'** |
|  | **'Right inferior frontal gyrus pars opercularis'** |
|  | **'Right precentral gyrus** |
